# Standing genetic variation as a major contributor to adaptation in the Virginia chicken lines selection experiment

**DOI:** 10.1101/018721

**Authors:** Zheya Sheng, Mats E. Pettersson, Christa F. Honaker, Paul B. Siegel, Örjan Carlborg

## Abstract

Artificial selection has, for decades, provided a powerful approach to study the genetics of adaptation. Using selective-sweep mapping, it is possible to identify genomic regions in populations where the allele-frequencies have diverged during selection. To avoid misleading signatures of selection, it is necessary to show that a sweep has an effect on the selected trait before it can be considered adaptive. Here, we confirm candidate selective-sweeps on a genome-wide scale in one of the longest, on-going bi-directional selection experiments in vertebrates, the Virginia high and low body-weight selected chicken lines. The candidate selective-sweeps represent standing genetic variants originating from the common base-population. Using a deep-intercross between the selected lines, 16 of 99 evaluated regions were confirmed to contain adaptive selective-sweeps based on their association with the selected trait, 56-day body-weight. Although individual additive effects were small, the fixation for alternative alleles in the high and low body-weight lines across these loci contributed at least 40% of the divergence between them and about half of the additive genetic variance present within and between the lines after 40 generations of selection. The genetic variance contributed by the sweeps corresponds to about 85% of the additive genetic variance of the base-population, illustrating that these loci were major contributors to the realised selection-response. Thus, the gradual, continued, long-term selection response in the Virginia lines was likely due to a considerable standing genetic variation in a highly polygenic genetic architecture in the base-population with contributions from a steady release of selectable genetic variation from new mutations and epistasis throughout the course of selection.

## Background

Adaptation is a dynamic evolutionary process where populations improve their fitness by accumulating beneficial alleles at loci controlling adaptive phenotypes. The polymorphisms contributing to adaptation can either be present as standing genetic variation at the onset of selection or emerge through mutations. A long-standing challenge in quantitative and evolutionary genetics has been quantification of the relative contributions from standing and emerging variation to long-term selection response [1, 2]. Although, such results are very difficult to obtain in studies of natural populations. Artificial selection provides an approach to study the origin and fate of beneficial mutations during adaptation [2].

Subjecting populations to artificial selection provides an accelerated evolutionary process that may result in extreme phenotypes with accompanying changes across the genome [2-4]. Using such experiments, the contribution by mutational variance to the evolution of quantitative traits can be quantified by, for example, measuring the release of genetic variance during selection from an inbred founder population [5]. Estimating the contribution from standing genetic variation to long-term selection response is, however, more complex. Whereas short-term contributions can be estimated based on the immediate selection response in a selection-experiment starting from outbred founders, long-term contributions of standing variation are more difficult to estimate because of confounding with effects of mutations that emerge over time. Other approaches are therefore needed to disentangle the contributions of standing variation from other sources of selectable additive variation such as new mutations or selection induced additive variation [6-9].

Here, we estimate the contribution of standing genetic variation to long-term selection-response in an experimental population developed by bi-directional selection from a common, segregating founder-population. This experimental design allows for the separation of contributions from standing genetic variation and new mutations even when molecular data on the base-population is missing [3, 4]. The key is to identify “soft sweeps” between the selected populations that result from selection on standing variation [10-13]. Here, the genetic effects of a large collection of such divergently fixed “soft” selective sweeps [3] were estimated to predict their contribution to selection response in the Virginia chicken lines. They provided insights into the dynamic processes involved in shaping the genetics of complex traits during adaptive evolution.

The base-population for the Virginia lines was established in 1957 by intercrossing 7 partially inbred lines of White Plymouth Rock chickens. The genetic variation entering the population thus represents a sample of the polymorphisms present when these partially inbred lines were founded. Since the founder population was established, the high (HWS) and low (LWS) body-weight lines have been bred with one new generation per year by single-trait, bi-directional selection for 56-day body-weight [14-16]. The response to selection has progressed steadily throughout the experiment, resulting in an 8-fold difference in 56-day body-weight after 40 generations of selection and currently, in the 57th generation, there is a 16-fold difference between the lines (Figure 1). Genome-wide comparisons between the divergently selected lines have identified more than 100 candidate “soft” adaptive sweeps between them [3, 4]. The contribution to selection response of these candidate selective-sweep regions that originate from the standing genetic variation is still unknown. Here, we identified which of these candidate selective-sweeps contributed to adaptation and estimated their individual and joint contributions to the adaptive trait. Further, because the confirmed “soft” sweeps originated from the standing genetic variation in the base-population, inferred will also be the amount of new selectable variation released during selection and then contributed to the long-term selection response.

**Figure 1:**
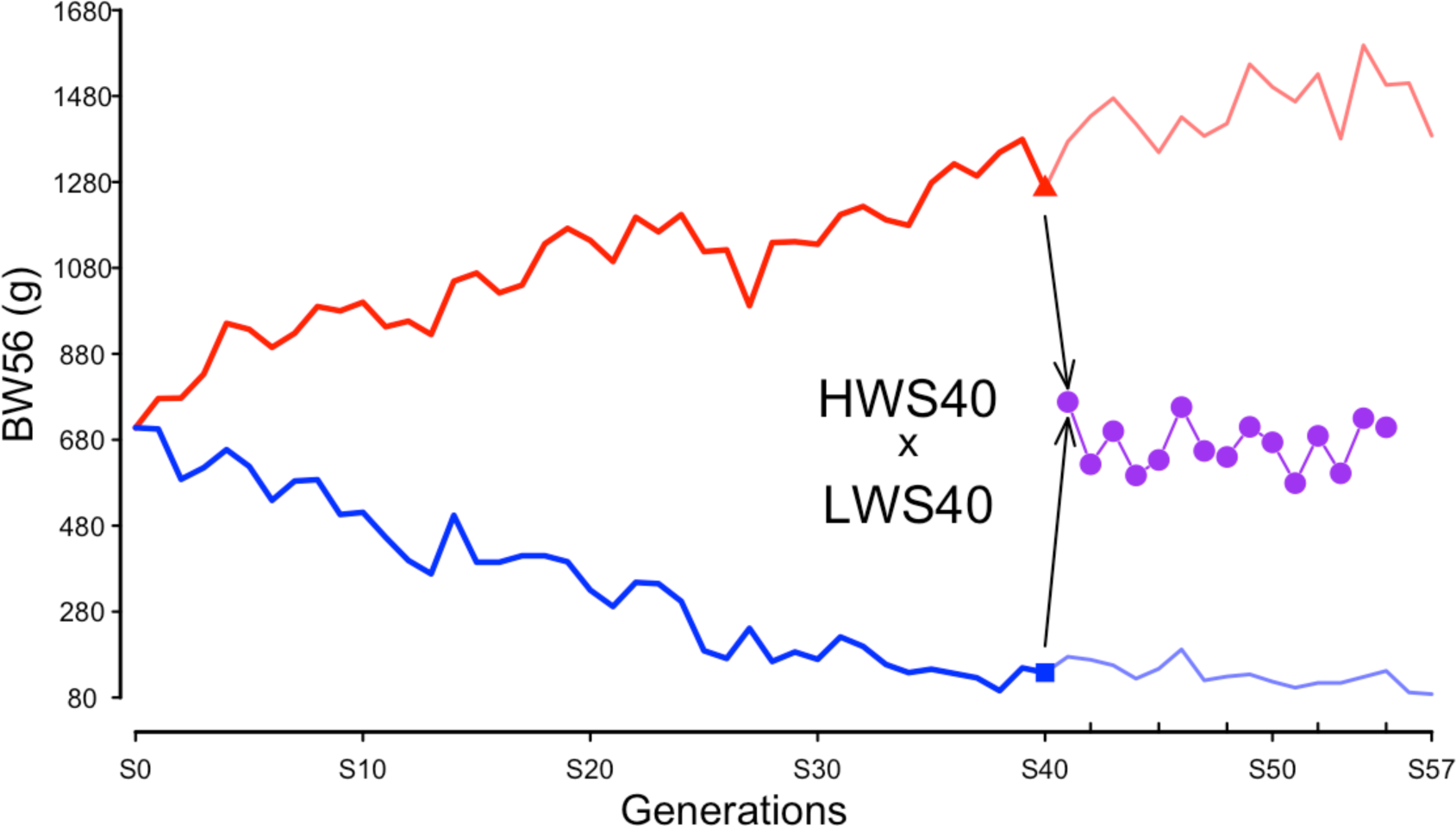
Body-weights at 56 days of age in the Virginia weight selected and Advanced Intercross Lines. Average body-weights per generation are provided for females in the high and low body-weight selected lines and as sex-averaged weights in the Advanced Intercross Line. BW56: 56-day body-weight.

This study describes a genome-wide approach to explore the contributions from a large number of selective-sweeps [3] to selection-response and adaptation in the Virginia chicken lines. A deep intercross population from HWS and LWS from generation S40, and genotyping and phenotyping a large F_15_ Advanced Intercross Line (AIL) generation for the selective-sweep regions, allowed us to estimate the contribution of these sweeps to the adaptive trait: 56-day body-weight. We found that the standing genetic variation at a large number of loci in the base-population was the major contributor to the selectable additive variance during adaptation. The highly polygenic genetic architecture for the selected trait in the base-population, together with a steady release of selectable genetic variation from new mutations and epistasis, provided a likely explanation for the gradual, continued, long-term response to selection in the Virginia lines.

## Results

Evaluated were the contribution by 106 selective-sweeps, where the high and low weight selected Virginia lines were fixed for alternative alleles, to selection response [3, 4]. Using genotypes for 252 markers in these sweeps, 99 clusters of markers that segregated independently were identified in the F_15_ generation of the AIL. In total, 38 of these regions were covered by a single marker and 61 regions by multiple markers. The physical length of the regions covered by multiple markers varied from 0.02 to 6.7 Mb and were distributed across most autosomes in the chicken genome (Figure 2; Additional file 1).

**Figure 2:**
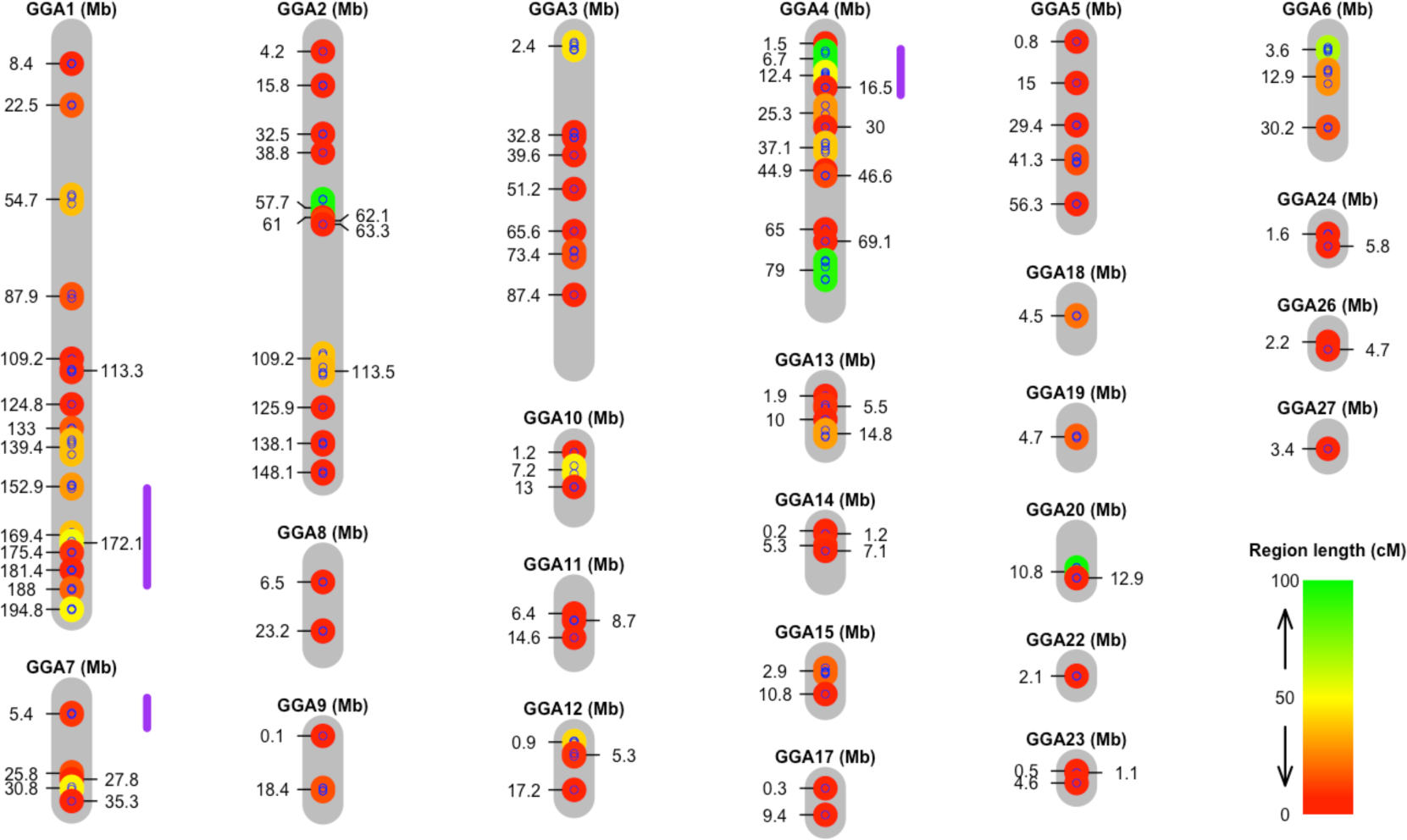
Genomic distribution of the selective sweeps. The gray bars represent chromosomes with their length in Mb on the Nov. 2011 (galGal4) genome assembly. The small blue dots indicate the locations of the 252 genotyped markers that passed quality control. The colored bars connecting the dots on the chromosomes illustrate the 99 independently segregating regions that were tested for association with 56-day body-weight in the F_15_ generation of the Virginia Advanced Intercross Line. Asterisks were used to indicate the presence of multiple sweeps in a region that could not be separated visually. The color of the bars indicate their genetic map lengths (cM, Haldane) in the F_15_ generation and the extension of their physical map lengths (Mb).

### Many adaptive selective-sweeps have contributed to selection-response

To estimate how many of the independently segregating regions contributed to 56-day body-weight in the AIL F_15_ generation, we initially selected one representative marker from each of the 99 targeted regions via a within-region, backward-elimination analysis. Then, an across-region analysis was performed to identify the set of regions that jointly contributed to the adaptive trait. A multi-locus, backward-elimination analysis was used where the final set of loci were selected at 5 and 20% False Discovery Rate (FDR) thresholds [17, 18]. The potential influences of population-structure were controlled using bootstrapping [19]. Table 1 summarizes the loci associated with 56-day body-weight at 5 and 20% FDR significance thresholds. Many of the 99 evaluated sweeps were associated with 56-day body-weight in the F_15_ AIL, 8 at a 5% and 16 at a 20% FDR. Thus, we confirmed that a large number of the candidate selective-sweeps identified previously [3] were adaptive selective-sweeps. Further, the standing genetic variation for 56-day weight in the base-population for the Virginia lines was highly polygenic, and considerable genetic variation originating from the standing variation has been exhausted after 40 generations of selection via the fixation within the divergent lines across these loci.

**Table 1.** Genetic effects of selective-sweeps associated with 56-day body-weight in the Virginia Advanced Intercross Line. Estimates are provided for one marker in each associated sweep. Genetic effects were estimated in generation F_15_ of the AIL.

| GGA<sup>1</sup> | Position<sup>2</sup><br>(Mb) | Marker | Additive<sup>3</sup><br>(a±SE) | Sign<sup>4</sup> | RMIP<sup>5</sup><br>(FDR 5/20%) | Sign<sup>6</sup> |
| --- | --- | --- | --- | --- | --- | --- |
| 1 | 87 | rs13899455 | -22.7±6.2 | 2.4×10<sup>-4</sup> | 0.63/0.71 | 5 % |
| 1 | 133 | rs13942473 | 16.5±8.0 | 3.9×10<sup>-2</sup> | 0.38/0.59 | 20 % |
| 1 | 142 | rs15448487 | 16.5±6.8 | 1.5×10<sup>-2</sup> | 0.37/0.65 | 20 % |
| 1 | 169 | rs14916997 | 28.9±6.3 | 6.0×10<sup>-6</sup> | 0.93/0.98 | 5 % |
| 2 | 61 | GGaluGA149337 | 25.8±7.9 | 1.2×10<sup>-3</sup> | 0.75/0.85 | 5 % |
| 2 | 112 | rs15143460 | 23.8±6.9 | 6.4×10<sup>-4</sup> | 0.55/0.66 | 5 % |
| 2 | 148 | rs15158686 | 19.0±6.3 | 2.8×10<sup>-3</sup> | 0.76/0.86 | 5 % |
| 3 | 34 | rs15321683 | 24.6±6.0 | 4.7×10<sup>-5</sup> | 0.65/0.73 | 5 % |
| 3 | 75 | GGaluGA228961 | 15.5±6.8 | 2.2×10<sup>-2</sup> | 0.28/0.47 | 20 % |
| 4 | 2 | rs14417942 | 14.5±6.7 | 3.0×10<sup>-2</sup> | 0.31/0.48 | 20 % |
| 4 | 12 | GGaluGA246087 | 17.8±6.6 | 7.3×10<sup>-3</sup> | 0.37/0.51 | 20 % |
| 4 | 45 | rs15560796 | 14.5±6.6 | 2.8×10<sup>-2</sup> | 0.39/0.55 | 20 % |
| 4 | 82 | rs14498744 | -31.3±6.6 | 2.0×10<sup>-6</sup> | 0.94/0.96 | 5 % |
| 10 | 9 | GGaluGA068581 | 11.3±6.4 | 8.0×10<sup>-2</sup> | 0.35/0.52 | 20 % |
| 13 | 10 | rs14059068 | 16.6±6.3 | 8.6×10<sup>-3</sup> | 0.42/0.61 | 20 % |
| 23 | 5 | rs15205573 | 18.5±6.6 | 5.4×10<sup>-3</sup> | 0.49/0.75 | 5 % |

<sup>1</sup>GGA: Gallus Gallus Autosome; <sup>2</sup> Nov. 2011 (galGal4) assembly; <sup>3</sup>Additive genetic effect ± Standard
Error estimated in model (C) or (D); <sup>4</sup>Significance for additive genetic effect in model including all loci
significant at 20% FDR; <sup>5</sup>RMIP: Resample Model Inclusion Probability [19] using 5 or 20% FDR
threshold [17]; <sup>6</sup>Significance thresholds 5/20% FDR that the marker was selected at with RMIP > 0.46
in after Bagging procedure [19] to correct for population structure

### The individual adaptive selective-sweeps have small allele-substitution effects

To estimate the contribution by the individual adaptive sweeps to the selected trait, additive, allele-substitution effects were estimated using a multi-locus association analysis. We found that the additive effects of the individual loci were generally small, and no individual locus had an allele-substitution effect greater than 29 g (or 0.2 σ_P_) for the selected trait (Table 1; Figure 3). The effects were similar for most of the loci when estimated in the F_15_ population (Figure 3A). Most of the HWS derived alleles increased weight, however, two regions also had transgressive effects on the trait - i.e. that an allele inherited from the LWS increased weight. The standing genetic variation in the base-population for the Virginia chicken lines thus has contributed with alleles of small effect across a highly polygenic genetic architecture.

**Figure 3:**
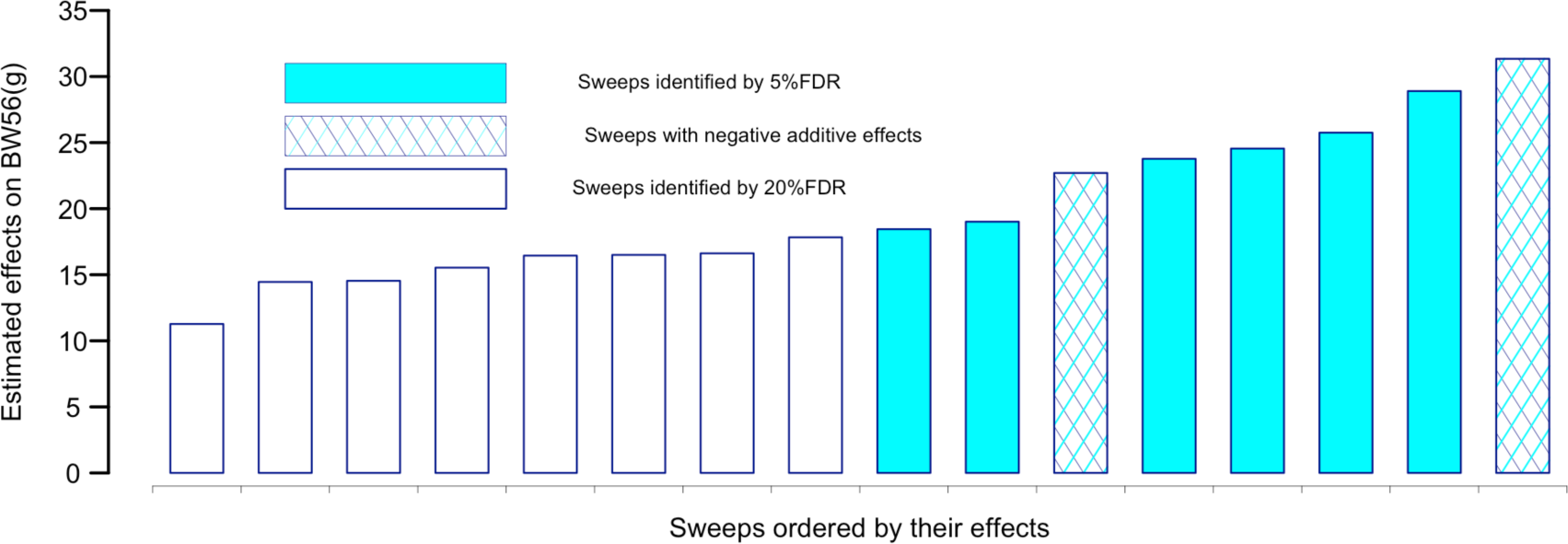
Allele-substitution effects of selective-sweeps associated with 56-day body-weight (BW56) in the Virginia Advanced Intercross Line. The effects were estimated in the F_15_ generation of the AIL. Colored bars indicate for selective-sweeps with associations at a 5% FDR and white bars selective-sweeps with associations at a 20% FDR. Solid colored bars indicate selective-sweeps where the HWS derived allele increases body-weight. Hashed colored bars indicate selective-sweeps where the LWS derived allele increase body-weight, i.e. regions that are transgressive.

### Large contribution by standing genetic variants to long-term selection response

When the joint contribution of the 16 adaptive selective-sweeps to the realised selection-response was estimated, none of the intercross chickens were HWS/HWS or LWS/LWS homozygous across all 16 regions. This provides a likely explanation for why the phenotypic range in the F_15_ intercross does not reach the high and low phenotypes of the HWS and LWS lines (Table 2). Therefore, we predict the joint contributions of the adaptive selective-sweeps to the divergence between the lines based on the average allele-substitution effect for HWS alleles across the selective-sweep regions shown to contribute to 56-day body-weight. The average estimates for the sweeps that were significant at 5/20% FDR were 15.7±2.5 (p = 6.7 × 10^-10^) /16.9±1.6 (p < 1 × 10^-16^) g, respectively. The predicted total contribution of these regions to the 1242 g line-difference between HWS40 and LWS40 (Table 2) are then 16 HWS alleles x 15.7 g = 251 g (20.2%) for the 5% set, and 32 HWS alleles × 16.9 g = 542 g (43.6%) for the 20% FDR set. Both of these estimates are biased downwards due to the inclusion of the two transgressive selective-sweep regions in the analysis. Thus, the total contribution by the adaptive selective-sweeps from the standing variation in the base-population to adaptation is likely to be at least 40% of the total realised response at generation 40.

**Table 2.** Summary statistics for the body-weights for the Advanced Intercross Line population. The AIL was bred between founders from generation 40 of the high (HWS) and low (LWS) selected lines.

| **Generation** | **BW56**<br>**(mean / $\sigma_P$)** |
| --- | --- |
| F<sub>0</sub> (HWS40) | 1412 / 125 |
| F<sub>0</sub> (LWS40) | 170 / 47 |
| F<sub>1</sub>-F<sub>15</sub> | 672 [569-756] / 148 [115-169] |

BW56: 56-day Body-Weight; $\sigma_P$ : Sex-averaged residual phenotypic standard deviation.

### Contribution by standing genetic variation to the selectable additive genetic variance

The next step was to estimate the contribution of the associated sweeps to the genetic variance in the F_15_ generation of the AIL. The narrow-sense heritability for 56-day weight in the F_15_, estimated based on the pedigree kinship [20], was 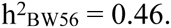. The 8 regions associated at the more stringent 5% FDR threshold together explained nearly one fifth of the residual phenotypic variance in 56-day body-weight (19.1% of residual 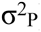). When we included the 8 additional regions selected at a 20% FDR, the estimate increased to 23.4% of 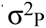. Thus, the associated selective-sweep regions contributed to approximately half of the additive genetic variance in this population (41.5 and 50.9% of 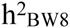 for regions significant at 5/20% FDR, respectively). Accordingly, the confirmed selective-sweeps have been major contributors to the selectable additive genetic variance during this selection experiment. Further, this result also illustrates that fixation at these adaptive selective-sweep regions depleted about half of the total additive genetic variance for 56-day body-weight during first the 40 generations of divergent selection.

### Relative contributions by standing genetic variation and other sources of variation to selection response

We estimated the amount of additive genetic variance present at onset of selection that was captured by the confirmed adaptive selective-sweeps by comparing the total additive genetic variation in the base-population obtained shortly after onset of selection [21] with the estimate of the additive genetic variance contributed by the adaptive selective-sweeps in the AIL F_15_ generation (Table 3). In total, the additive variance of the sweeps in the AIL corresponded to 70 to 86% of the total additive variance in the base-population. This result suggests that the adaptive selective-sweeps identified and confirmed here represented a major part of the standing genetic variation in the base-population. It also implies that other sources of variation, such as novel mutations and selection induced genetic variation, have contributed to the gradual long-term selection response in the Virginia chicken lines.

**Table 3.**
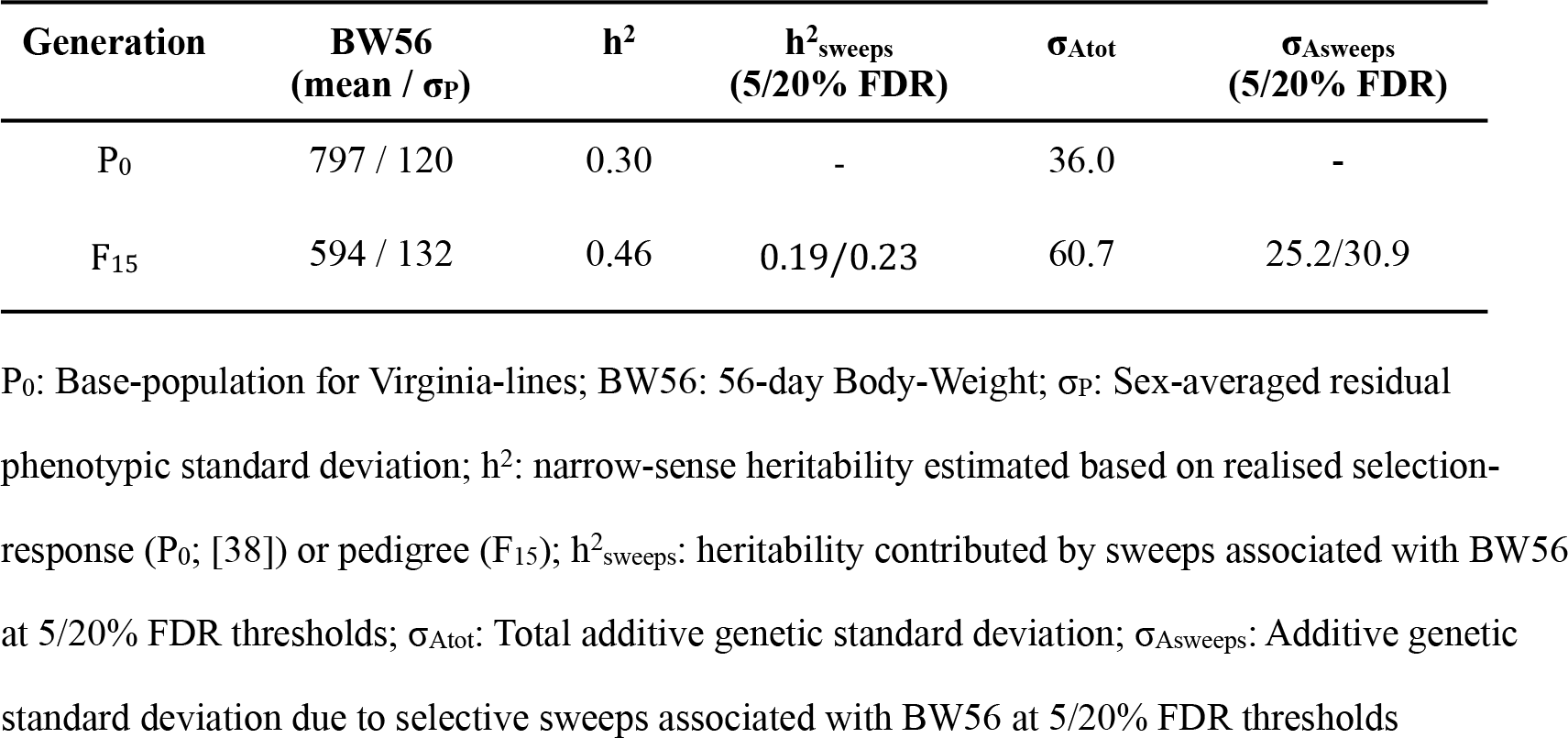
Summary of estimates of genetic parameters in the Virginia lines. Estimates are provided for the base-population of the Virginia chicken lines (P_0_) and the F_15_ generation from the Advanced Intercross Line population (F_15_).

### Discussion

#### Many selective-sweeps associated with 56-day body-weight

In total, 8/16 candidate selective-sweep regions were associated with 56-day body-weight at 5/20% FDR. Most of the loci selected in the 5% FDR set should be true positive associations as < 1 false positive is expected at this significance level, whereas a few false-positive associations could be present among the loci selected at 20% FDR.

In the original selective-sweep mapping study [3], simulations predicted that no less than 40%, but most likely considerably more than half of the candidate selective-sweeps originating from the base-population would result from selection rather than drift. In light of results reported here, this theoretical prediction still appears realistic. First, we confirmed that the genetic-architecture of 56-day weight in the base-population was highly polygenic and that the contributions by individual loci were small. Also, quantitative genetics estimates were in agreement with this because they also predicted that the confirmed sweeps did not represent all of the standing additive genetic variance in the base-population. As the power to detect loci with small individual effects was limited in the association analysis of only 800 F_15_ individuals, it is likely several loci with similar or smaller effects than those confirmed here contributed to the trait but remained undetected. Second, although the current study covered 99 candidate selective-sweeps, it was only a sample of all candidate selective-sweeps present in the Virginia-lines. Because some candidate sweeps would have been missed in the study [3], which was based on medium-density genotyping from a 60k SNP chip rather than whole-genome re-sequencing. Furthermore, some of the candidate selective-sweep regions previously reported [3] were missing from our dataset due to failed genotyping, whereas the effects of others could not be separated due to the resolution in the AIL F_15_ generation. To obtain a complete understanding of the genetic architecture of selection-response due to standing variation in the base-population of the Virginia chicken lines will require additional candidate selective-sweeps.

#### Base-population alleles contribute to selection-response via small individual genetic effects

The individual allele-substitution effects for the confirmed adaptive selective-sweep regions were small (Figure 3). No individual locus had an allele-substitution effect greater than 29 g (0.2 residual σ_P_ ; Figure 3; Table 1) corresponding to a contribution of less than 3% of the residual 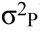 despite the average minor-allele frequencies being rather high (average MAF = 0.21).

The base-population for the Virginia selection experiment involved intercrossing seven moderately inbred lines of White Plymouth Rock chickens [14-16]. During inbreeding, alleles with negative effects on fitness are likely to be removed from a population. From earlier work, it is known that several of the major growth-promoting alleles that have emerged during domestic animal improvement programs also have negative pleiotropic effects on fitness. Extreme examples include the susceptibility to Malignant hypothermia by the Ryanodine receptor mutation that promotes growth in pigs [22] and the calving problems due to muscle-growth promoting Myostatin mutations in cattle [23]. Our finding that most alleles from the base-population have small effects on growth is consistent with the presence of a negative correlation between enhanced or reduced growth and fitness.

That most alleles originating from the standing genetic variation had small effects also follows the expectation from theoretical studies involving soft sweeps. The probability of fixation for alleles with small effects is higher when selection acts on standing genetic variation than on a new mutation as there is a higher likelihood that a weakly selected new mutation will be lost [12].

#### The alleles originating from the standing genetic variation in the base-population make a large joint contribution to the additive variance and selection-response

The confirmed adaptive sweeps contributed about 40% of the realised selection-response and half of the total additive genetic variance in the F_15_ population. A significant portion of the phenotypic divergence between the lines was thus due to alleles originating from the standing genetic variation in the base-population and for which the lines are now fixed for alternative alleles. This clearly illustrates the importance of the standing genetic variation in the base-population for the gradual, long-term response to selection observed throughout this experiment. Some of this can be explained by additional loci originating from the base-population that either had individual effects that were too small to be detected in the population-size available for this study, were located on candidate selective-sweeps that were fixed at generation 40 but not included in this study, or were in regions not yet fixed for alternative alleles at generation S40.

Two of the confirmed selective-sweeps were transgressive, i.e. the allele inherited from the low-weight selected line increased weight in the F_15_ generation. This confirms the theoretical work by Robertson, who in 1960 showed that alleles of small effect can be fixed by chance in the opposite direction to that of selection [24]. Similar findings have also been reported in studies of other artificially selected populations (see e.g. [25, 26]).

Overall, our findings are consistent with those reported for another long-term selection experiment: the Illinois corn selection lines [25, 27], in particular the highly polygenic genetic architectures with many loci of small individual effects as well as the presence of transgressive loci. This suggests that the selection on highly polygenic genetic architectures, where many loci make minor contributions rather than on emerging mutations with large effects, has been important for the long-term success of these selection experiments. Such studies will continue to deliver valuable insights not only to the general features of polygenic architectures contributing to morphological traits in vertebrates, but also the basic processes involved in accelerated evolution and adaptation.

#### Long-term selection response from the joint contributions of standing genetic variation, novel mutations and epistasis

The adaptive selective sweeps in the F_15_ generation of the AIL explained from 42 to 51% of the total additive genetic variance in this intercross, and 70 to 86% of the total additive genetic variance present for 56-day body-weight in the base-population (Table 3). Much of the standing genetic variation has been exhausted while producing an 8-fold difference in the selected trait during the first 40 generations of selection. This supports the theoretical expectation that standing variation will be an important contributor to the initial selection response for a population subjected to a novel selection pressure [28]. However, because standing variation was not exhausted even after an intense artificial long-term, single-trait selection, this suggests an adaptive value for standing variation over longer periods of time. This may be especially relevant in natural populations where selection is not as intense.

Further, we estimated that approximately 40% of the total selectable additive genetic variation that contributed to the long-term selection response emerged throughout the experiment from other sources of variation. Some contributions are expected from new mutations [28], however, contributions may involve other sources such as selection induced genetic variation [6]. Although we have not explicitly explored the contributions by these here, previous reports involving the Virginia lines have provided insights on this topic. For example, a major contribution has been reported from a network of interacting loci that via a capacitating epistatic mechanism was likely to have induced considerable selectable additive variation in response to selection [7-9, 29]. Moreover, a recent study of the within-line response to selection has also identified a novel allele that due to its rapid fixation within the high-weight selected line was likely a newly arisen major allele [4]. Further work is, however, needed to quantify these sources of variation in relation to that of the standing variation.

## Conclusions

Here, we empirically confirm that the gradual response to long-term, bi-directional, single-trait selection in the Virginia chicken lines was, to a great extent, due to standing genetic variation across a highly polygenic genetic architecture in the common base-population. A large number of loci, each having small allele-substitution effects, were major contributors to the selectable additive variance during adaptation. Much of the variation from the base-population was fixed in the divergent lines after 40 generations of selection. The considerably larger total additive genetic variance present within and between the lines after 40 generations of selection suggests that important contributions have also been made by a steady release of selectable genetic variation from new mutations and epistasis. In summary, these results provide not only novel insights to the genetics contributing to the gradual, continued, long-term response to selection in the Virginia lines, but also to the fundamental genetic mechanisms contributing to selection and adaptation.

## Methods

### Ethics statement

All procedures involving animals used in this experiment were carried out in accordance with the Virginia Tech Animal Care and Use Committee protocols.

### Animals and phenotyping

The animals used in this study were from an Advanced Intercross Line (AIL) bred from generation S40 parents from two lines of chickens divergently selected for juvenile body-weight: the Virginia high (HWS) and low (LWS) lines. The HWS and LWS lines were founded in 1957 from a base-population obtained by crossing seven partially inbred lines of White Plymouth Rock chickens. Since then, they have been subjected to bi-directional selection for a single trait, high or low 56-day body-weight (BW56), respectively, and currently the lines have reached generation 58 (Figure 1). More detailed information on the selected populations is available [14-16].

A complete description of the development of the Advanced Intercross population can be found in the publications describing the analysis of data from the F_2_-F_8_ generations of the AIL [29, 30]. The AIL (Figure 1) was founded by F_0_ parents from generation S40 of the HWS and LWS lines, whose sex-average BW56 at that generation were 1412 g (SE: ± 36 g) and 170 g (SE: ± 5 g), respectively. In total, 907 individuals were hatched in generation F_15_. Out of these, 852 survived until 56 days of age, when their body-weight was measured. More details on the phenotypes of the founder lines and the AIL are provided in Table 2.

### DNA extraction

Blood samples were collected using a sterile needle and syringe, and then transferred to a tube containing disodium EDTA. DNA was extracted from whole blood samples using a Qiagen DNeasy kit.

### Marker selection and genotyping

In an earlier study, Johansson *et al.* [3] identified a large number of SNPs that were fixed for alternative alleles across the genome of the Virginia lines based on 60K SNP chip genotypes obtained for individuals from generations 40 and 50 of the HWS and LWS lines. These SNPs were clustered into selective-sweep regions: 116 clusters containing 998 SNPs in generation 40 and 163 clusters with 1746 SNPs in generation 50. We selected 316 SNP markers to cover the 134 autosomal selective-sweep regions. As some markers were selected for fixation in generation 50, all were not fixed in the founders for the intercross obtained from generation 40. They were, however, all highly informative with an allele frequency difference ≥ 0.9 between LWS and HWS. Samples from all F_15_ birds with recorded 56-day weights (n=852) were sent for genotyping at the 316 selected SNPs using the GoldenGate assay (Illumina, CA) at the SNP technology platform in Uppsala (Sweden).

In total, 27 individuals and 64 SNPs failed to fulfill one or more of the following quality control criteria: call rate of individuals >0.9, call frequency of SNPs >0.9, minor allele frequency >0.05. They were removed from the subsequent analysis, resulting in a final dataset consisting of genotypes for 252 SNPs in 825 birds. In total, the failed genotyping resulted in a loss of 26 initially targeted regions, resulting in a final coverage for 106 regions [3], 72 of which were present in both generation 40 and 50, and 34 emerged at generation 50.

### Reclustering and summaries of targeted selective sweep regions

After ordering the genotyped markers based on their physical locations on the galGal4 assembly, we redefined the clusters based on their genetic linkage in the F_15_ generation. This was to define clusters that segregated independently in the F_15_ generation in order to identify the number of independent, targeted selective-sweeps that contribute to 56-day body-weight in the lines. The criteria used for clustering markers were to assign adjacent markers that were no more than 50cM apart into the same cluster and limit the range of each cluster to cover no more than 100cM. According to these criteria, 99 independent regions were defined for use in the association analyses. The linkage between the markers in cM (Haldane mapping function) was estimated using the function est.map from the R/qtl package in R [31] where the function for an inbred intercross population could be used as the markers in this study are fixed, or nearly fixed, in the analysed population. Although a small number of the genotyped markers were not fixed for alternative alleles in the F_0_ founders, we decided that the obtained estimates of linkage were sufficiently precise for re-clustering of the markers.

In total, we then screened for associations between 252 markers that passed quality control and 56-day body-weight in 825 chickens from the F_15_ generation of the Advanced Intercross Line (AIL). To estimate how many of the independently segregating regions contributed to 56-day body-weight in the AIL F_15_ generation, we initially selected one representative marker from each of the 99 independently segregating clusters of markers, as defined above, in the F_15_ generation (Figure 2; Additional file 1) via a within-region, backward-elimination analysis. As mentioned above, these markers were located in 106 of the selective-sweeps regions detected in the genome in two previous studies [3, 4].

### Heritability estimation

To estimate the heritability in the narrow sense for 56-day body-weight, we used a linear mixed model:

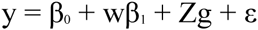

where y is the phenotype of 56-day body-weight, *β*_*0*_ is an intercept term, *β*_*1*_ is the sex effect and *w* the associated indicator vector. Furthermore, the random polygenic effects *g* are normally distributed with correlation matrix given by the pedigree-kinship matrix *A*, i.e. 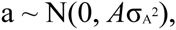, *Z* is the associated incidence matrix, and *ε* is a normally distributed residual error 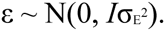. The pedigree kinship matrix *A* was constructed using the R/pedigree package [32]. The linear mixed model was fitted using the R/hglm package [33] and the heritability estimated as 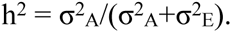.

### Association analyses

#### Defining line-origin for the genotyped marker-alleles

The GoldenGate genotyping assay report F_15_ genotypes in an “ATCG” basis without information about the line-origin of the respective alleles. Therefore, we first transformed the genotypes into a “-1 0 1” basis by comparing the genotypes in the F_15_ to that of the HWS/LWS F_0_ founders. “-1” was used to represent the case when both alleles were of LWS line origin, “0” that the individual was heterozygous, and “1” that both alleles were of HWS line-origin. In this way, positive estimates of the additive effects in the association analysis indicate that the HWS derived allele increased weight, and negative additive effects that the LWS derived allele increased weight (i.e. that it is transgressive).

### Selection of markers to represent the independently segregating selective-sweep regions

The dataset was not sufficiently large to separate the effects of the linked markers within the 99 selective-sweep regions. Therefore, the first step in the association analysis was to simplify the subsequent analyses by selecting one marker in each of these regions to represent their joint effects in the subsequent multi-locus analysis. To select these markers, we used a within-locus backward-elimination approach in a linear model framework. The additive genetic effects of all markers in the selective-sweep region to be evaluated were then included together with the fixed effects of the sex for the bird. The analysis was thus based on the following model:

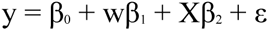

where *y* is the phenotype, *β*_*0*_ is an intercept term, *β*_*1*_ is the sex effect and *w* the associated indicator vector, *β*_*2*_ is the set of additive sweep effects modeled as fixed effects and *X* is the associated design matrix coded as -1, 0, 1 for the line origin of the marker genotypes, and *ε* is a normally distributed residual error. The number of markers within each region was limited and no problem of confounding between the fixed and random effects was detected.

Using backward-elimination from the full model, we then identified the individual marker within each region that had the most significant effect, without requiring a particular significance for it at this stage of the analysis. This analysis was performed using custom written scripts in R.

### A multi-locus, backward-elimination analysis to identify adaptive selective-sweep regions

The objective of the confirmation-study was to identify the set of selective-sweep regions that jointly contribute to 56-day body-weight in the F_15_ generation of the AIL. The statistical analysis was chosen with the background knowledge that the genetic architecture of body-weight in this population was highly polygenic [3, 4, 30, 34] and that potentially as many as half of the genotyped selective-sweep regions contributed to weight [3]. As all individuals in the AIL were progeny of dams of the same age, hatched on the same date and reared separate from their parents, environmental contributions to between-family means in the F_15_ population may be considered minimal. Thus, a large portion of the difference in family means should be due to the joint effects of the many selective-sweep regions studied. When a large multi-locus mixed model was fitted to the data and fixed effects of markers across multiple selective-sweep regions were included together with a polygenic random effect in the model to account for family-effects, there was a strong confounding between the fixed and random effects. This confounding may be explained by an assumption of linear mixed models being violated. A linear mixed model y = Xβ + Zu + e assumes that there is no correlation between a column in *X*, or a linear combination of several columns, with the true random effect *u*. This might occur where the number of columns in *X* is large, and thus we deemed this analysis as unsuitable for use in the multi-locus analysis. However, as population-structure might be of concern [35, 36], we chose to validate our results using a bootstrap-based approach developed for the same purpose for general deep-intercross populations, including Advanced Intercross Lines, by Valdar *et al.* [19]. These bootstrap analyses were implemented in custom scripts in the statistical software R [37].

The bootstrap based approach was implemented in a backward-elimination model-selection framework across the genotyped selective-sweeps. Before doing the multi-locus analysis, we evaluated whether it was statistically appropriate to perform this analysis across all the 99 regions that segregated independently in the linkage analysis. For this, a standard measure to identify potential high-order collinearity, the “variance inflation factor” or VIF, was used. Consistent with the linkage analysis, there was no large pair-wise correlation between the 99 markers. However, some marker genotypes could (almost) be written as a linear combination of the genotypes at all the other markers which would lead to collinearity problems in a multi-locus statistical analysis. Therefore the markers for the affected selective-sweeps (21, 22, 80, 89, 90) were removed from the subsequent analyses.

The true size of the model (i.e. the number of contributing regions) is unknown in advance and could range substantially from nearly half of the tested regions to a few individual associations. To compare models with such a wide range of variables to include, we opted to use an adaptive model selection criterion controlling the False Discovery Rate [17, 18] developed for this purpose. The multi-locus models used during backward-elimination were implemented in a standard linear model framework, starting with a full model including the fixed effects of sex and the additive effects of the 99 selected markers from the independently segregating regions. Convergence was based on two alternative FDR levels: 5% and 20%. The analysis was performed both in the original data, and using bootstrapping with 1000 resamples. For the bootstrap analyses, the RMIP (Resample Model Inclusion Probability) was calculated for all regions in backward-elimination analyses with 5 and 20% FDR among the selected loci as termination criteria. A final model was decided for each FDR level by selecting the regions with RMIP > 0.46 as suggested for an AIL generation F_18_ [19]. The additive genetic effects for each locus was estimated using the multi-locus genetic model described below including regions selected at 5 and 20% FDR.

To produce Figure 3, the allele substitution effects were estimated for all markers selected at 20% FDR. This was done in a multi-locus linear-model framework, where the genotypes of the markers in other significant regions selected in the same analysis were included as co-factors. The residual variances explained by the selected regions at 5 and 20% FDR were computed as the proportion of the residual variance of the null-model including only the sex of the individual.

### Predicting the fraction of the HWS40-LWS40 line-difference explained by the associated selective-sweep regions

Two alternative approaches were used to predict how much of the population difference of 1242 g between the parental HWS and LWS from generation 40 (Table 2) could be explained by the selected regions at 5% or 20% FDR. This was done by regressing the individual’s phenotype to the total number of alleles of HWS origin carried across the evaluated regions. Using this approach, the standard-error of the estimate for the average allele-substitution effect was lower than for the individual estimates of the loci. This analysis was performed across the regions selected at 5 and 20% FDR. The obtained estimates were then multiplied by the total number of allele-substitutions between the HWS and LWS lines for this set of loci to obtain an estimate of the contribution to the line-difference.

## List of abbreviations

HWS: Virginia <u>H</u>igh Body-<u>W</u>eight <u>S</u>elected line; LWS Virginia <u>L</u>ow Body-<u>W</u>eight <u>S</u>elected line; BW56: Chicken <u>B</u>ody-<u>W</u>eight at <u>56</u> days of age; AIL: <u>A</u>dvanced <u>I</u>ntercross <u>L</u>ine; GGA: <u>G</u>allus <u>G</u>allus <u>A</u>utosome

## Authors’ contributions

ÖC and PBS initiated the study and designed the project with MEP; PBS developed the Virginia chicken lines; PBS designed, planned and bred the Virginia Advanced Intercross Line; PBS and CFH designed, planned, bred, bled, phenotyped, and extracted DNA from the F_15_ Virginia Advanced Intercross Line population; MEP was responsible for selecting polymorphisms in the selective-sweep regions, developing primers for these and sending samples for genotyping; ZS performed the quality control of the genotype data; ÖC designed the statistical analyses; ZS and ÖC contributed analysis scripts; ÖC and ZS performed the data analyses, summarized the results, and wrote the manuscript. All authors read and approved the final manuscript.

## Acknowledgements

We thank Leif Andersson for initiating the intercross experiment with PBS and sharing the QTL mapping data from the F_2_ intercross. Stefan Marklund and Andreas Lundberg are acknowledged for their valuable contributions during selection of polymorphisms in the selective-sweep regions for genotyping, development of primers for these, and for interacting with the genotyping centre. We also thank Lars RÖnnegård for useful input on the statistical analysis and the manuscript. Genotyping was performed by the SNP&SEQ Technology Platform in Uppsala, which is part of Science for Life Laboratory at Uppsala University and is supported as a national infrastructure by the Swedish Research Council (VR-RFI).

---

**Additional Table 1.**
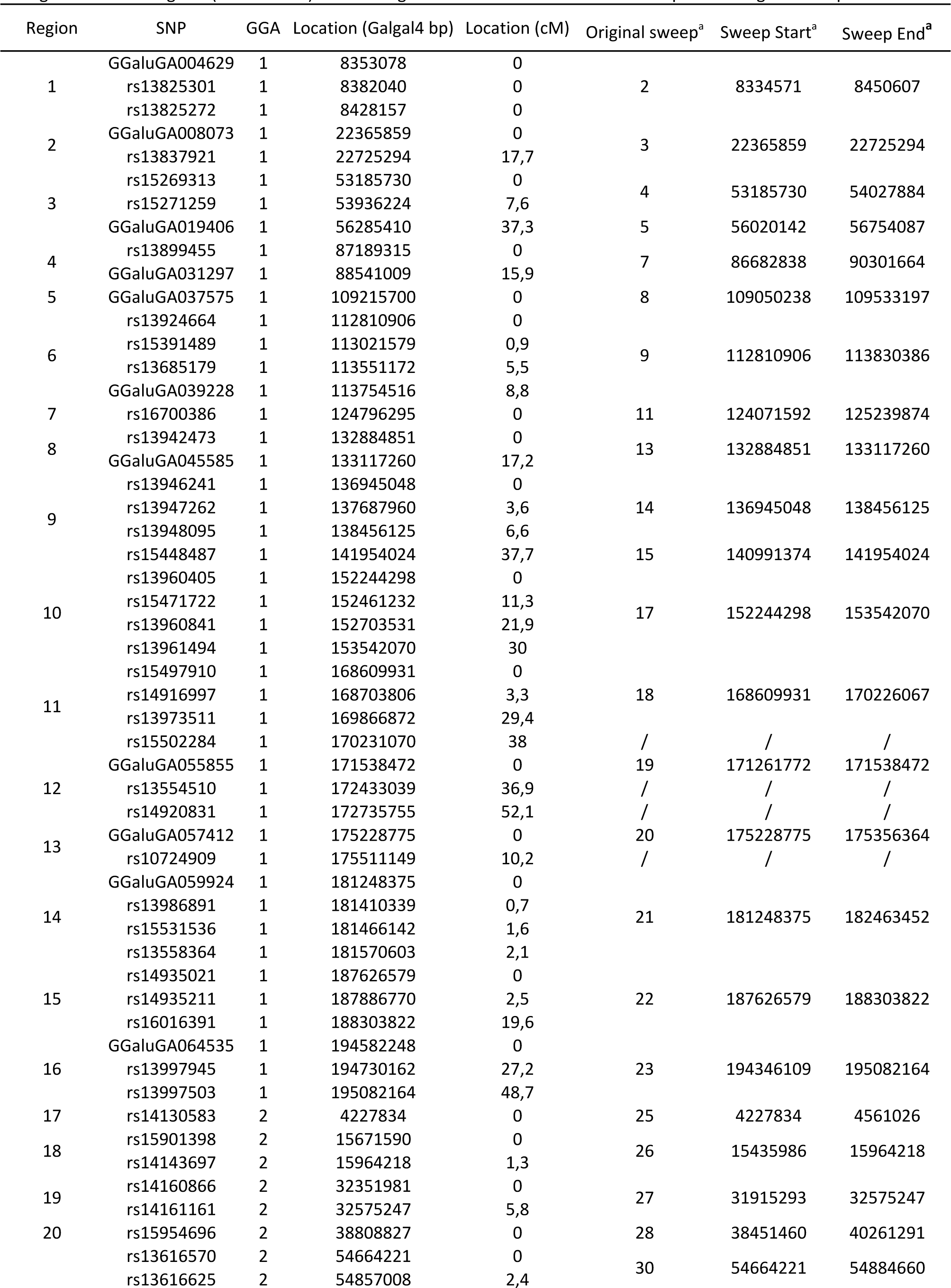
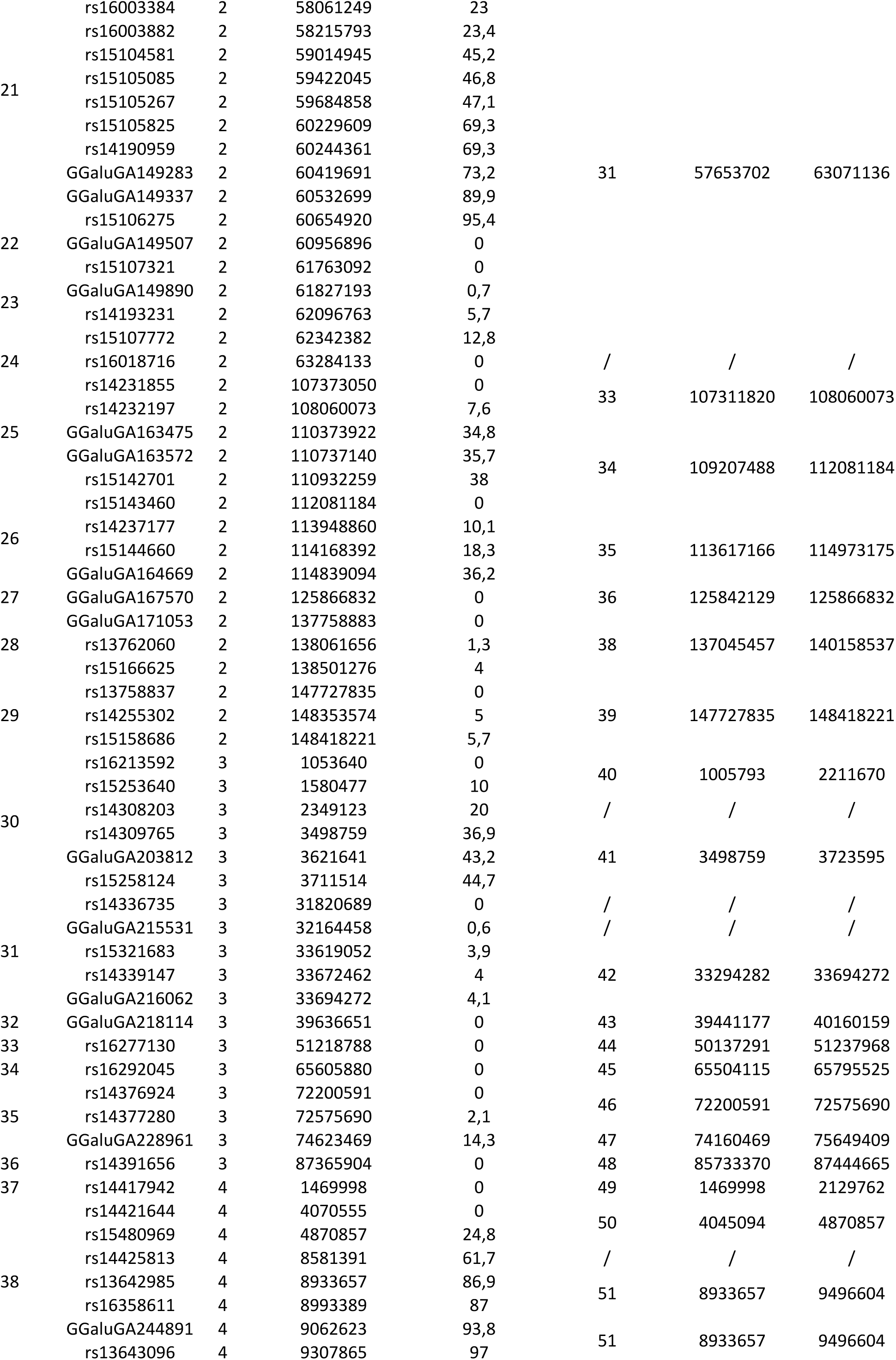
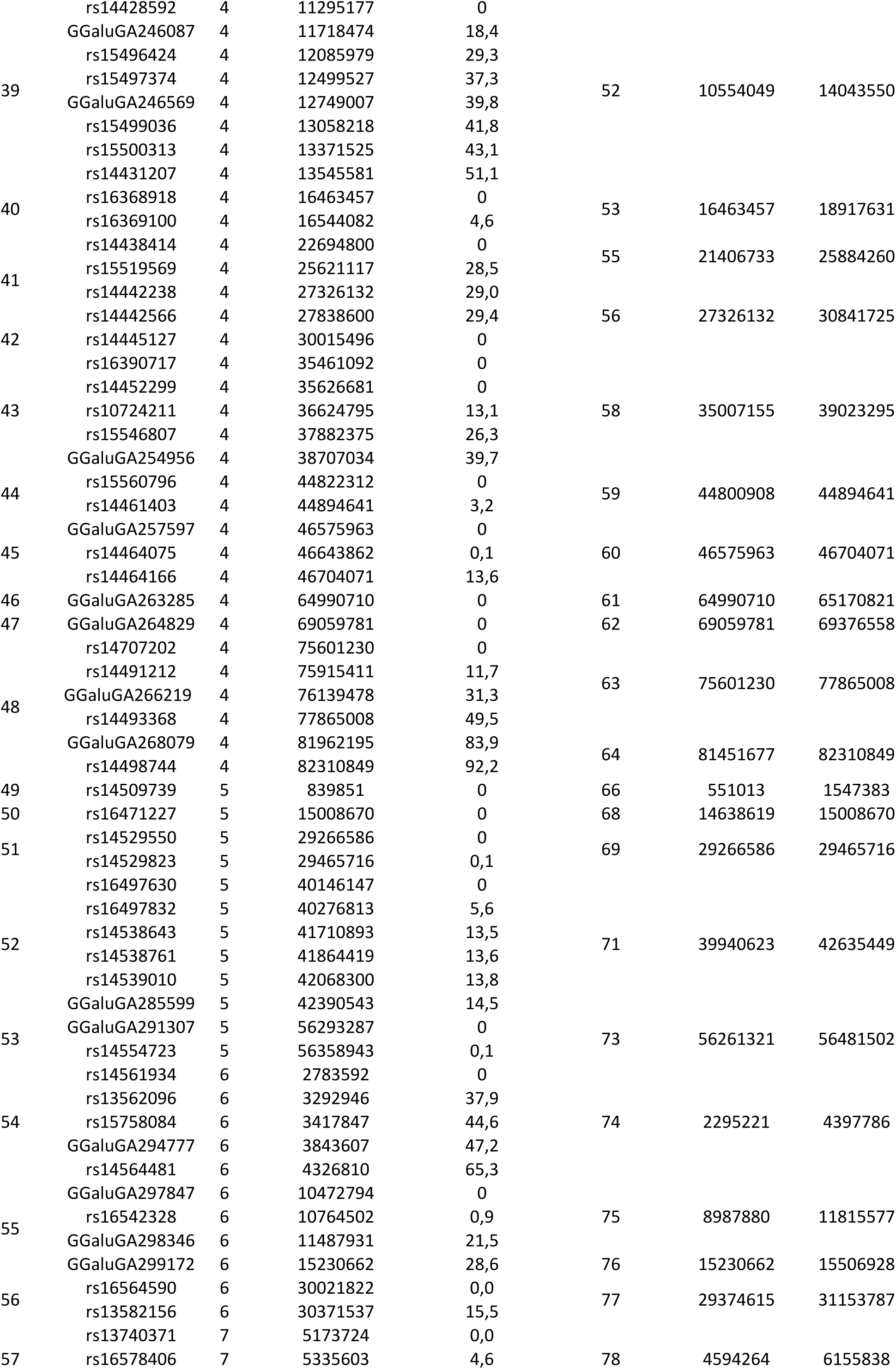
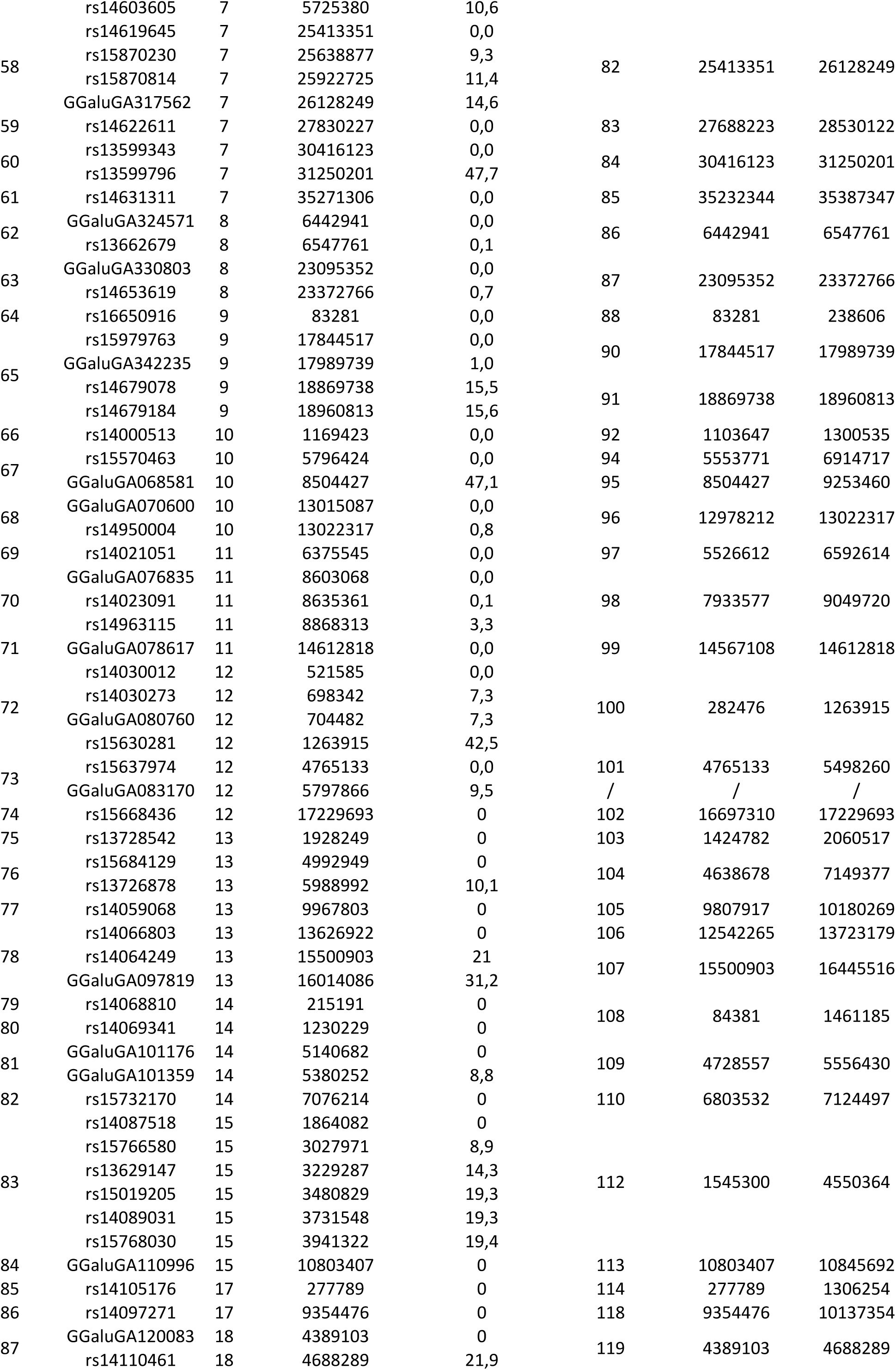
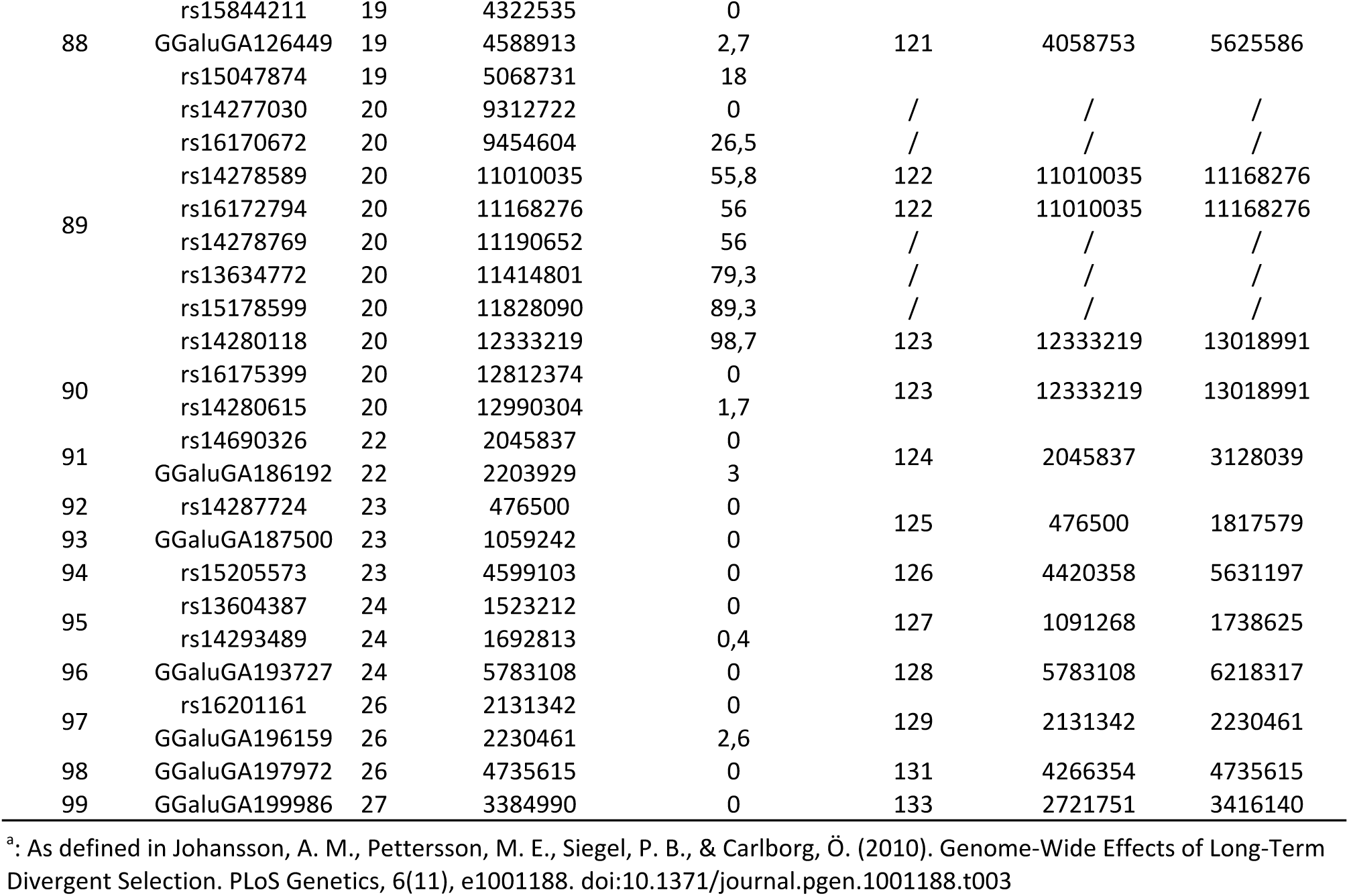
The 99 independently segregating regions in the F_15_ and the 252 SNPs passing QualityEControl within them. Information is provided regarding both their physical location on the galGal4 genomeEassembly (bp) and their linkages within the regions (cM Haldane) in the F_15_ generation as well as the relationship to the original sweeps^a^.

## References

1. Orr HA: The population genetics of beneficial mutations. Philosophical Transactions of the Royal Society B: Biological Sciences 2010, 365:1195–1201.

2. Burke MK: How does adaptation sweep through the genome? Insights from long-term selection experiments. Proceedings of the Royal Society B: Biological Sciences 2012, 279:5029–5038.

3. Johansson AM, Pettersson ME, Siegel PB, Carlborg Ö: Genome-Wide Effects of Long-Term Divergent Selection. PLoS Genet 2010, 6:e1001188.

4. Pettersson ME, Johansson AM, Siegel PB, Carlborg Ö: Dynamics of Adaptive Alleles in Divergently Selected Body Weight Lines of Chickens. G3 (Bethesda) 2013, 3:2305–2312.

5. Hill WG, Caballero A: Artificial selection experiments. Annual Review of Ecology and Systematics 1992, 23:287–310.

6. Eitan Y, Soller M: Selection Induced Genetic Variation. In Evolutionary Theory and Processes: Modern Horizons. Edited by Wasser S. Dordrecht: Springer Netherlands; 2004:153–176.

7. Carlborg Ö, Jacobsson L, Åhgren P, Siegel P, Andersson L: Epistasis and the release of genetic variation during long-term selection. Nat Genet 2006, 38:418–420.

8. Le Rouzic A, Siegel PB, Carlborg Ö: Phenotypic evolution from genetic polymorphisms in a radial network architecture. BMC Biol 2007, 5:50.

9. Le Rouzic A, Carlborg Ö: Evolutionary potential of hidden genetic variation. Trends in Ecology & Evolution 2008, 23:33–37.

10. Orr HA, Betancourt AJ: Haldane’s sieve and adaptation from the standing genetic variation. Genetics 2001, 157:875–884.

11. Przeworski M, Coop G, Wall JD: The signature of positive selection on standing genetic variation. Evolution 2005, 59:2312–2323.

12. Hermisson J, Pennings PS: Soft sweeps: molecular population genetics of adaptation from standing genetic variation. Genetics 2005, 169:2335–2352.

13. Pennings PS, Hermisson J: Soft sweeps II--molecular population genetics of adaptation from recurrent mutation or migration. Mol Biol Evol 2006, 23:1076–1084.

14. Dunnington EA, Siegel PB: Long-term divergent selection for eight-week body weight in white Plymouth rock chickens. Poultry science 1996, 75:1168–1179.

15. Márquez GC, Siegel PB, Lewis RM: Genetic diversity and population structure in lines of chickens divergently selected for high and low 8-week body weight. Poultry science 2010, 89:2580–2588.

16. Dunnington EA, Honaker CF, McGilliard ML, Siegel PB: Phenotypic responses of chickens to long-term, bidirectional selection for juvenile body weight-historical perspective. Poultry science 2013, 92:1724–1734.

17. Abramovich F, Benjamini Y, Donoho DL, Johnstone IM: Special Invited Lecture: Adapting to Unknown Sparsity by Controlling the False Discovery Rate. The Annals of Statistics 2006, 34:584–653.

18. Gavrilov Y, Benjamini Y, Sarkar SK: An Adaptive Step-Down Procedure with Proven Fdr Control Under Independence. The Annals of Statistics 2009, 37:619–629.

19. Valdar W, Holmes CC, Mott R, Flint J: Mapping in structured populations by resample model averaging. Genetics 2009, 182:1263–1277.

20. Sinnwell JP, Therneau TM, Schaid DJ: The kinship2 R package for pedigree data. Hum Hered 2014, 78:91–93.

21. Siegel P: A double selection experiment for body weight and breast angle at eight weeks of age in chickens. Genetics 1962, 47:1313–1319.

22. Fujii J, Otsu K, Zorzato F, de Leon S, Khanna VK, Weiler JE, O’Brien PJ, MacLennan DH: Identification of a mutation in porcine ryanodine receptor associated with malignant hyperthermia. Science 1991, 253:448–451.

23. Grobet L, Martin LJ, Poncelet D, Pirottin D, Brouwers B, Riquet J, Schoeberlein A, Dunner S, Ménissier F, Massabanda J, Fries R, Hanset R, Georges M: A deletion in the bovine myostatin gene causes the double-muscled phenotype in cattle. Nat Genet 1997, 17:71–74.

24. Robertson A: A Theory of Limits in Artificial Selection. Proceedings of the Royal Society B: Biological Sciences 1960, 153:234–249.

25. Laurie CC, Chasalow SD, LeDeaux JR, McCarroll R, Bush D, Hauge B, Lai C, Clark D, Rocheford TR, Dudley JW: The genetic architecture of response to long-term artificial selection for oil concentration in the maize kernel. Genetics 2004, 168:2141–2155.

26. Parts L, Cubillos FA, Warringer J, Jain K, Salinas F, Bumpstead SJ, Molin M, Zia A, Simpson JT, Quail MA, Moses A, Louis EJ, Durbin R, Liti G: Revealing the genetic structure of a trait by sequencing a population under selection. Genome Research 2011, 21:1131–1138.

27. Hill W: A century of corn selection. Science 2005, 307:683–684.

28. Barrett RDH, Schluter D: Adaptation from standing genetic variation. Trends in Ecology & Evolution 2008, 23:38–44.

29. Pettersson M, Besnier F, Siegel PB, Carlborg Ö: Replication and explorations of high-order epistasis using a large advanced intercross line pedigree. PLoS Genet 2011, 7:e1002180.

30. Besnier F, Wahlberg P, Rönnegård L, Ek W, Andersson L, Siegel PB, Carlborg Ö: Fine mapping and replication of QTL in outbred chicken advanced intercross lines. Genet Sel Evol 2011, 43:3.

31. Broman KW, Wu H, Sen S, Churchill GA: R/qtl: QTL mapping in experimental crosses. Bioinformatics 2003, 19:889–890.

32. pedigree: Pedigree functions [http://CRAN.R-project.org/package=pedigree]. Accessed 29 Apr 2015.

33. Rönnegård L, Shen X, Alam M: hglm: A package for fitting hierarchical generalized linear models. The R Journal 2010, 2:20–28.

34. Jacobsson L, Park H-B, Wahlberg P, Fredriksson R, Perez-Enciso M, Siegel PB, Andersson L: Many QTLs with minor additive effects are associated with a large difference in growth between two selection lines in chickens. Genet Res 2005, 86:115–125.

35. Peirce JL, Broman KW, Lu L, Chesler EJ, Zhou G, Airey DC, Birmingham AE, Williams RW: Genome Reshuffling for Advanced Intercross Permutation (GRAIP): Simulation and Permutation for Advanced Intercross Population Analysis. PLoS ONE 2008, 3:e1977.

36. Cheng R, Lim JE, Samocha KE, Sokoloff G, Abney M, Skol AD, Palmer AA: Genome-Wide Association Studies and the Problem of Relatedness Among Advanced Intercross Lines and Other Highly Recombinant Populations. Genetics 2010, 185:1033–1044.

37. R Development Core Team: R: a Language and Environment for Statistical Computing. Vienna, Austria; 2015. Accessed 29 Apr 2015.

38. Siegel PB: Selection for Body Weight at Eight Weeks of Age: 1. Short term Response and Heritabilities. Poultry science 1962, 41:954–962.

